# Bile Salts and N-Acetyl-D-Glucosamine induce transcription of the *vieSAB* Operon in *Vibrio cholerae* El Tor

**DOI:** 10.1101/2022.02.02.478790

**Authors:** Lauren M. Shull, Andrew Camilli

**Affiliations:** Department of Molecular Biology and Microbiology, Graduate School of Biomedical Sciences, Tufts University School of Medicine, Boston, Massachusetts, USA

## Abstract

The *vieSAB* operon regulates motility, biofilm formation and cholera toxin production in the classical biotype of *V. cholerae*. Classical biotype mutants lacking the cyclic-di-GMP phosphodiesterase and response regulator *vieA* or its cognate sensor kinase *vieS* fail to effectively colonize the infant mouse small intestine, are less motile than the wild-type strain, and are hyper-biofilm formers. However, deletion of *vieA* or *vieS* in the currently-circulating El Tor biotype does not have a demonstrable effect on these phenotypes. To begin to understand the role of *vieSAB* in the El Tor biotype, we studied its regulation and the effect of *vieA* expression on bacterial physiology, separating the functional roles of VieA’s enzymatic and DNA-binding activities. We identify bile salts and N-acetylglucosamine, which can be found at high concentration in the small intestine, as stimulating expression of *vieSAB* during *in vitro* growth. We show that, in contrast to many response regulators, VieA does not regulate its own expression. Together, these data indicate that *V. cholerae* upregulates *vieSAB* in response to small-intestinal signals, allowing VieA to alter the physiology of the cell through its transcriptional regulation and cyclic-di-GMP signaling activities in order to adapt to the human host.

## Introduction

*Vibrio cholerae*, the causative agent of cholera, is classified into serogroups according to the specific makeup of the O-antigen on its lipopolysaccharide (LPS). Of the over 140 serogroups, the O1 is the predominant causes of cholera (Chatterjee and Chaudhuri, 2003). O1 is further divided into serotypes Inaba and Ogawa which differ by a terminal methyl group (Chatterjee and Chaudhuri, 2003). O1 is also divided along a different axis into two biotypes, classical and El Tor. The classical biotype was responsible for causing cholera pandemics until the beginning of the seventh pandemic in 1961 (Banerjee et al., 2014). The El Tor biotype is causing the still ongoing seventh pandemic and has completely supplanted the classical biotype as a cause of cholera (Shahid et al., 1984). The El Tor biotype is more easily transmissible but causes less-severe disease, two factors that contributed to its spread and its ability to outcompete the classical biotype (Shahid et al., 1984). Genetically, classical and El Tor biotypes differ by hundreds of single-nucleotide polymorphisms and by alleles associated with the cholera toxin-encoding CTX prophage (Faruque et al., 2007).

Cyclic-di-GMP was first described as an activator of cellulose synthesis in *Gluconacetobacter xylinus* (formerly *Acetobacter xylinum*) (Ross et al., 1986; Ross et al., 1987; Ross et al., 1990). It was later shown to function as a second messenger in classical biotype *V. cholerae*, where it positively regulates biofilm formation (Tischler and Camilli, 2004). C-di-GMP is synthesized by enzymes called diguanylate cyclases (DGCs), which are characterized by GGDEF domains (Ryjenkov et al., 2005). Proteins with either EAL or HD-GYP domains are capable of catalyzing the degradation of c-di-GMP and both belong to a class of enzymes called phosphodiesterases (PDEs). It is not uncommon for single proteins to incorporate both a GGDEF domain and a PDE domain.

The role of c-di-GMP in pathogenic bacteria is complex. In general, a high concentration of c-di-GMP results in biofilm formation, and a low concentration of c-di-GMP results in virulence factor expression and motility. These phenotypes were initially described in classical biotype *V. cholerae* following the discovery of an EAL-domain PDE, VieA (Tischler and Camilli, 2004; Tischler and Camilli, 2005; Tamayo et al., 2005). Transcriptional responses to varying c-di-GMP concentrations in *V. cholerae* corroborate the phenotypic findings: high levels of c-di-GMP result in decreased expression of flagellar genes and increased expression of *Vibrio* exopolysaccharide (*vps*) genes which synthesize the exopolysaccharide that makes up the biofilm extracellular matrix (Beyhan et al., 2006). *V. cholerae* encodes at least 31 PDEs and 44 DGCs, whose activity can respond to many environmental signals to alter the overall concentration of c- di-GMP in the cell (Galperin et al., 2001; Conner et al., 2017). In the El Tor biotype, the responses to changes in c-di-GMP concentration do not match the responses noted in the classical biotype (Tamayo et al., 2008). It is likely that different DGCs and PDEs are active in each biotype and that the interplay of the many DGCs and PDEs results in the differing behaviors that have been noted.

### The *vieSAB* operon of *V. cholerae* encodes a three-component phosphorelay system

VieS and VieA function as a two-component system in which VieS is a sensor kinase and VieA is its cognate response regulator (Tischler and Camilli, 2002). The environmental signal sensed by VieS remains unknown. VieS specifically phosphorylates VieA at the aspartic acid residue in position 52, the characteristic phosphorylation residue of response regulators (Martinez-Wilson et al., 2008). In addition to the EAL domain which confers PDE activity, VieA includes a helix-turn-helix DNA-binding domain characteristic of response regulators. The third protein, VieB, is expressed only during infection and has a negative regulatory role on VieS and VieA activity by dampening the phosphorelay (Mitchell et al., 2015).

The *vieSAB* operon was also identified as a regulator of cholera toxin production. In classical biotype *V. cholerae*, mutants of *vieS* or *vieA* have colonization defects in the infant mouse model of infection (Tischler and Camilli, 2002). In the classical biotype, deletion mutants of *vieA* also have reduced motility and form excessive biofilm (Tischler and Camilli, 2004; Tischler and Camilli, 2005; Martinez-Wilson et al., 2008). These phenotypes of the *vieA* mutant are largely due to dysregulated c-di-GMP levels in the absence of VieA’s PDE activity (Martinez-Wilson et al., 2008). *In vitro* assays showed that VieS can specifically phosphorylate VieA and that phosphorylation of VieA enhances its activity, for which the hypothesized mechanism is positive autoregulation as phosphorylation does not directly enhance the enzymatic activity of the EAL domain, and autoregulation is common in two-component system response regulators (Martinez-Wilson et al., 2008; Tamayo et al., 2005).

In contrast to the classical biotype, the *vieSAB* operon is dispensable for infection by the El Tor biotype (Tamayo et al., 2008). The *vieSAB* operon is also expressed at a lower level in the El Tor biotype during growth in laboratory conditions (Beyhan et al., 2006). As in the classical biotype, however, the *vieSAB* operon in the El Tor biotype is expressed during animal infection (Mandlik et al., 2011). The differences in expression of the *vieSAB* operon likely contribute to its differential role in classical versus El Tor biotype. Here we sought to expand our understanding of how *vieSAB* is regulated in the El Tor biotype.

## Materials and Methods

All strains and plasmids used are listed in Table 1. The O1 El Tor wave three *V. cholerae* strain HC1037 was used throughout unless otherwise specified. Unless otherwise specified, bacterial cultures were grown at 37 °C in Miller lysogeny broth (LB; BD Difco), in M9 chemically-defined minimal medium (BD Difco), or on solid agar plates consisting of LB solidified with 1.5% agar. Liquid cultures were aerated by rolling or shaking. Antibiotics for selection and propagation of *V. cholerae* and/or *Escherichia coli* were used at the following concentrations: streptomycin 100 μg/mL, kanamycin 50 μg/mL, chloramphenicol 10 μg/mL, ampicillin 100 μg/mL. Arabinose induction was carried out using a concentration of L-arabinose of 1 mg/mL.

**Table 1.** Bacterial strains and plasmids used in this study.

| Strain | Description | Reference |
| --- | --- | --- |
| WT | *V. cholerae* O1 El Tor Ogawa isolate from Haiti 2014 | Duncan et al., 2018 |
| *E. coli* BL21(DE3) | *E. coli* BL21(DE3), a wild-type *E. coli* especially well-suited for expression from P<sub>BAD</sub> promoters | Wood, 1966 |
| WT pBAD18-Kan *vieA* | WT containing pBAD18-Kan with wild-type *vieA* controlled by an arabinose-inducible promoter | This study |
| WT pBAD18-Kan *vieA*<sup>D52E</sup> | WT containing pBAD18-Kan with a phosphomimetic *vieA* mutation (D52E) controlled by an arabinose-inducible promoter | This study |
| WT pBAD18-Kan *toxT* | WT containing pBAD18-Kan with wild-type *toxT* controlled by an arabinose-inducible promoter | This study |
| WT pBBR-lux p<sub>*vieSAB*</sub> | WT containing plasmid pBBR-lux with *luxCDABE* under the control of the *vieSAB* promoter | This study |
| WT pBBR-lux p<sub>*tcpA*</sub> | WT containing plasmid pBBR-lux with *luxCDABE* under the control of the *tcpA* promoter | This study |
| WT *vieA* + p<sub>*vieSAB*</sub> | WT containing both pBAD18-Kan *vieA* and pBBR-lux p<sub>*vieSAB*</sub> | This study |
| WT *vieA*<sup>D52E</sup> + p<sub>*vieSAB*</sub> | WT containing both pBAD18-Kan *vieA*<sup>D52E</sup> and pBBR-lux p<sub>*vieSAB*</sub> | This study |
| WT *toxT* + p<sub>*vieSAB*</sub> | WT containing both pBAD18-Kan *toxT* and pBBR-lux p<sub>*vieSAB*</sub> | This study |
| WT *vieA* + p<sub>*tcpA*</sub> | WT containing both pBAD18-Kan *vieA* and pBBR-lux p<sub>*tcpA*</sub> | This study |
| WT *vieA*<sup>D52E</sup> + p<sub>*tcpA*</sub> | WT containing both pBAD18-Kan *vieA*<sup>D52E</sup> and pBBR-lux p<sub>*tcpA*</sub> | This study |
| WT *toxT* + p<sub>*tcpA*</sub> | WT containing both pBAD18-Kan *toxT* and pBBR-lux p<sub>*tcpA*</sub> | This study |
| *E. coli* pBAD18-Kan *vieA* | *E. coli* BL21(DE3) containing pBAD18-Kan with wild-type *vieA* controlled by an arabinose-inducible promoter | This study |
| *E. coli* pBAD18-Kan *vieA*<sup>D52E</sup> | *E. coli* BL21(DE3) containing pBAD18-Kan with a phosphomimetic *vieA* mutation (D52E) controlled by an arabinose-inducible promoter | This study |
| *E. coli* pBAD18-Kan *toxT* | *E. coli* BL21(DE3) containing pBAD18-Kan with wild-type *toxT* controlled by an arabinose-inducible promoter | This study |
| *E. coli* pBBR-lux<br>p<sub>vieSAB</sub> | *E. coli* BL21(DE3) containing plasmid pBBR-lux with *luxCDABE* under the control of the *vieSAB* promoter | This study |
| *E. coli* pBBR-lux p<sub>tcpA</sub> | *E. coli* BL21(DE3) containing plasmid pBBR-lux with *luxCDABE* under the control of the *tcpA* promoter | This study |
| *E. coli* *vieA* +<br>p<sub>vieSAB</sub> | *E. coli* BL21(DE3) containing both pBAD18-Kan *vieA* and pBBR-lux p<sub>vieSAB</sub> | This study |
| *E. coli* *vieA*<sup>D52E</sup> +<br>p<sub>vieSAB</sub> | *E. coli* BL21(DE3) containing both pBAD18-Kan *vieA*<sup>D52E</sup> and pBBR-lux p<sub>vieSAB</sub> | This study |
| *E. coli* *toxT* +<br>p<sub>vieSAB</sub> | *E. coli* BL21(DE3) containing both pBAD18-Kan *toxT* and pBBR-lux p<sub>vieSAB</sub> | This study |
| *E. coli* *vieA* +<br>p<sub>tcpA</sub> | *E. coli* BL21(DE3) containing both pBAD18-Kan *vieA* and pBBR-lux p<sub>tcpA</sub> | This study |
| *E. coli* *vieA*<sup>D52E</sup> +<br>p<sub>tcpA</sub> | *E. coli* BL21(DE3) containing both pBAD18-Kan *vieA*<sup>D52E</sup> and pBBR-lux p<sub>tcpA</sub> | This study |
| *E. coli* *toxT* +<br>p<sub>tcpA</sub> | *E. coli* BL21(DE3) containing both pBAD18-Kan *toxT* and pBBR-lux p<sub>tcpA</sub> | This study |
| *V. cholerae* *vieSAB-bla-neo* | *V. cholerae* E7946 with promoterless *bla* (Ap<sup>R</sup>) and *neo* (Km<sup>R</sup>) inserted immediately downstream of the native *vieSAB* locus | This study |
| **Plasmid** | **Description** | **Reference** |
| pBAD18-Kan | pBAD-derivative plasmid encoding kanamycin resistance on the backbone and a pBR322 origin | Guzman et al., 1995 |
| pBBR-lux | Mobilizable plasmid encoding *V. harveyi* *luxCDABE*, *cat*, and a pBBR1 origin | Lenz et al., 2004 |

### Generation of mutants

Mutations in *V. cholerae* were generated by cotransformation as described in Dalia et al., 2014. *V. cholerae* El Tor strains harboring plasmids were similarly generated by natural transformation. *E. coli* strains harboring plasmids were generated by electroporation at 1.8 kV in 1 mm cuvettes. Primers used in this study are listed in Table 2. Variants of pBBR-lux and pBAD18-Kan were constructed using traditional restriction endonuclease molecular cloning.

**Table 2:**
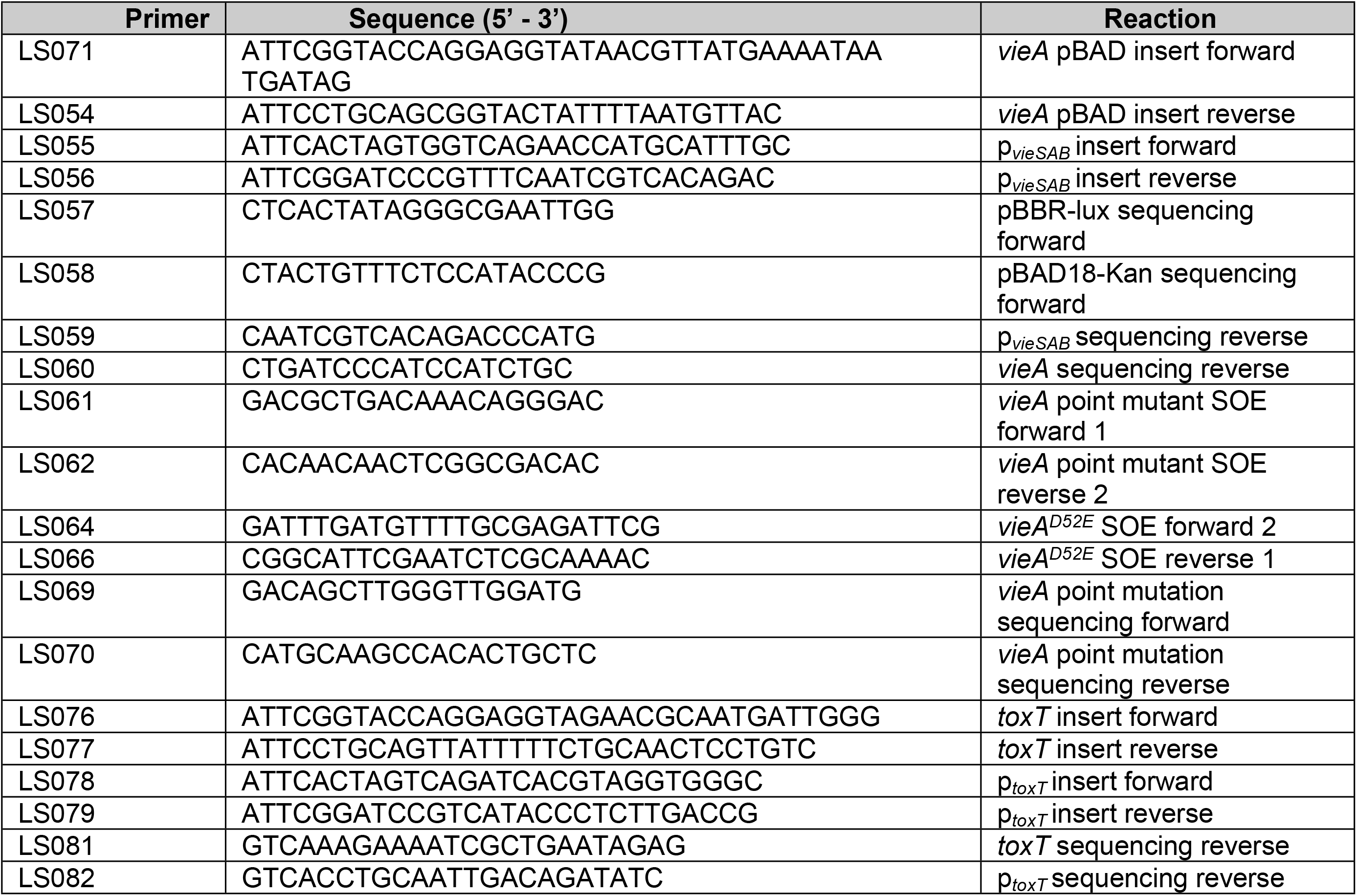
Primers used in this study.

Single-base alterations to plasmid sequences were performed using a PCR technique based on QuikChange site-directed mutagenesis (Stratagene).

### Dual-plasmid and luciferase assays

Dual-plasmid assays consist of bacterial strains containing both pBAD18-Kan with a transcription factor of interest inserted under the control of the arabinose-inducible promoter P_BAD_, and pBBR-lux with a promoter of interest inserted to control the transcription of *luxCDABE*. Strains containing these plasmids were grown overnight and then diluted into clear-bottomed white-sided 96-well plates (Corning) using 100 μL LB medium, and incubated at 37 °C with shaking in a Biotek Synergy H1 plate reader with the lid on. Each well was measured for OD_600_ and luminescence periodically over the course of the experiment.

### Electrophoretic mobility-shift assays

Electrophoretic mobility-shift assays (EMSA) were performed using native 6% polyacrylamide gels cast and run in 1X tris-borate-EDTA buffer (TBE). Purified His-VieA was used from a prior study (Mitchell et al., 2015). Fluorescent DNA probes were generated by PCR including dCTP-Cy5. Primers used for probe generation are listed in Table 2. Probes and purified protein were incubated in the presence of sheared calf-thymus DNA and BSA for 15 minutes at 30 °C in the dark before being loaded onto the polyacrylamide gel. Polyacrylamide gel electrophoresis (PAGE) was carried out at 4 °C for approximately two hours using 150 V.

### Antibiotic resistance reporter assays

Antibiotic resistance assays were used to measure transcription from the native *vieSAB* promoter in different growth conditions. This assay used a strain of El Tor *V. cholerae* with insertions of promoterless genes for ampicillin resistance (*bla*) and kanamycin resistance (*neo*) directly following *vieB* at its native locus. A suitable ribosome-binding site was provided for each reporter gene. When transcription of *vieSAB* is low, the reporter strain remains sensitive to concentrations of both antibiotics as low as 5 μg/mL. However, in growth conditions were transcription of *vieSAB* is high, the reporter strain is capable of growing in up to 100 μg/mL of each antibiotic. To screen for conditions that induce expression of *vieSAB*, a mid-exponential phase LB culture of this reporter strain was pelleted, washed with PBS, and finally resuspended in PBS. This washed culture was quantified by plating for CFU and used to inoculate six cultures in the condition of interest; bacteria were allowed to grow to mid-exponential phase for three hours at 37 °C with shaking unless otherwise noted. Ampicillin and kanamycin were added to three of the cultures at a concentration of 50 μg/mL and all six cultures were allowed to grow for one additional hour. Finally, bacterial survival was quantified by plating for CFU.

## Results

### VieA does not directly regulate expression of *vieSAB*

We developed a dual-plasmid luciferase reporter technique to investigate whether VieA carried out positive autoregulation, like many other two-component system response regulators. On plasmid pBAD18-Kan we can induce expression of VieA; a phosphomimetic VieAD52E which we hypothesize will have enhanced DNA-binding activity; or ToxT, a protein whose regulatory activity is well-understood and which can serve as a positive control when paired with its target promoter. On a second plasmid based on pBBR-lux, the luciferase operon of *V. harveyi*, *luxCDABE*, is under the control of either the *vieSAB* promoter P_*vieSAB*_ or the *tcpA* promoter P_*tcpA*_. The protein ToxT is known to positively regulate its own expression via the upstream promoter P_*tcpA*_ so this pair of plasmids acts as a positive control. Mismatched pairs of plasmids such as the VieA expression plasmid with the P_*tcpA*_ reporter plasmid were used as negative controls.

We generated *V. cholerae* strains harboring each plasmid individually, and all pairwise combinations. We monitored OD_600_ and light production in relative light units (RLU) from all strains in the presence or absence of the inducer, arabinose (Fig 1A). The positive control, *toxT* expression induced by arabinose paired with the *tcpA* promoter driving luciferase expression, shows robust light production as expected. Light production drops as cells approach stationary phase, which is expected as luciferase expression and light production is metabolically costly and available energy decreases when cells enter stationary phase. The conditions of interest, VieA or VieAD52E expressed with arabinose and paired with the *vieSAB* promoter driving luciferase expression, show no light production. This suggests that in the context of wild-type *V. cholerae* El Tor, VieA does not directly bind the *vieSAB* promoter and positively regulate expression.

**Fig 1.**
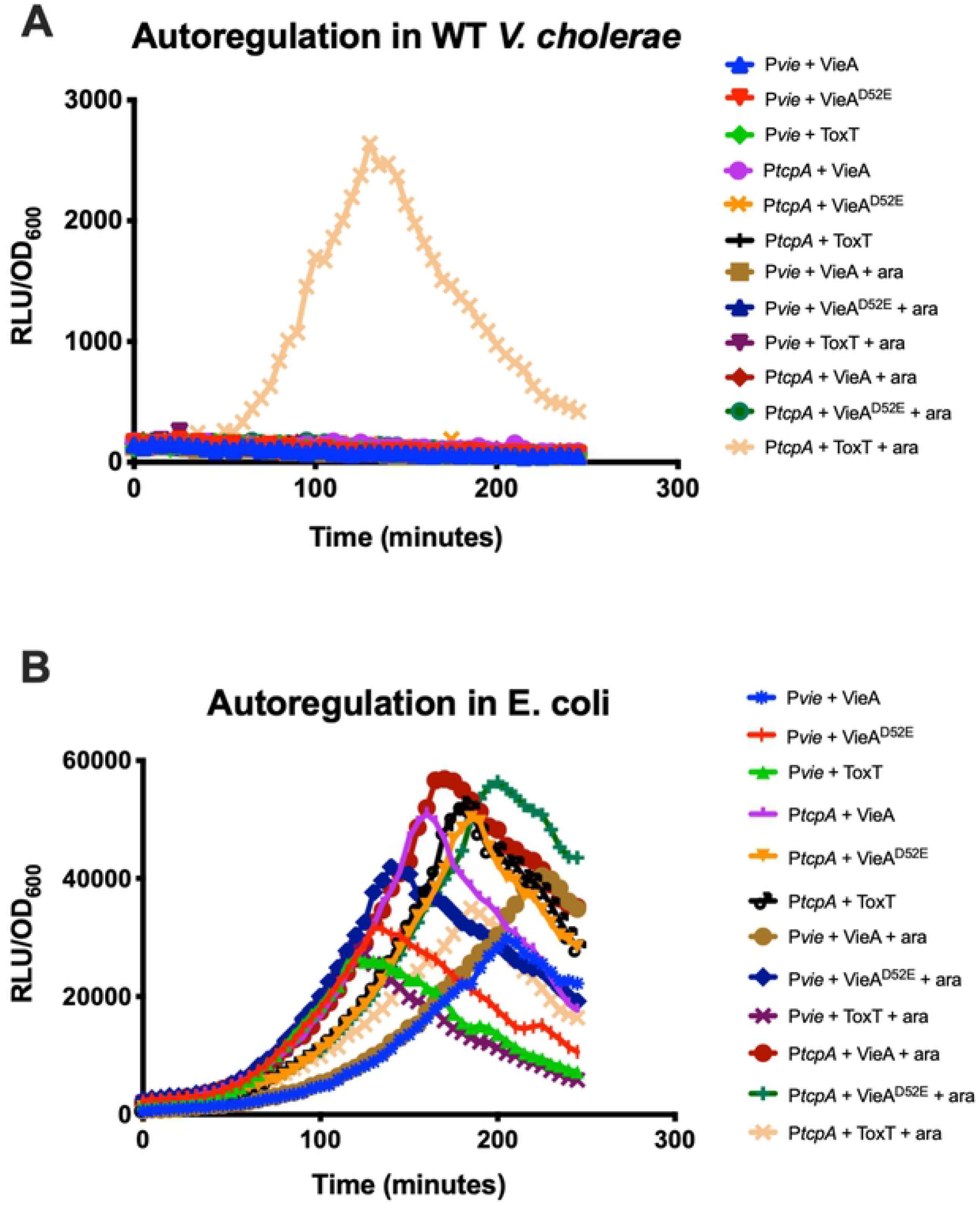
Reporter assay for *vieSAB* promoter activity. Autoregulation of the *vieSAB* promoter by VieA measured by dual-plasmid luciferase assay in A) *V. cholerae* or B) heterologous host *E. coli*.

In order to isolate the potential VieA-P_*vieSAB*_ interaction from other regulation occurring in *V. cholerae*, we reconstructed the same combinations of plasmids in the heterologous system of *E. coli*. All strains harboring each pairwise combination of expression and reporter plasmids were generated in *E. coli*, and the same luciferase assay carried out (Fig 1B). There is general derepression of the *vieSAB* and *tcpA* promoters in *E. coli*, as all strains containing a pBBR-lux derived plasmid produced luminescence regardless of the paired pBAD-derived plasmid and regardless of induction by arabinose.

Though we did not observe positive autoregulation by VieA, it was still possible that VieA could be acting as a repressor at the *vieSAB* promoter, or that it can bind but is unable to overcome repression mediated by other factors in *V. cholerae*. To test whether VieA can bind its own promoter, we carried out an EMSA in which we incubated His_6_-VieA with a fluorescent DNA probe representing the entire intergenic region between *vieS* and the upstream gene; this is the same region which was cloned into pBBR-lux to drive expression of the luciferase operon. We added the high-energy phosphate donor acetyl phosphate in order to phosphorylate VieA, as phosphorylation may be required for DNA binding. We did not observe any shift in mobility of the DNA probe (S1 Fig), indicating VieA does not by itself bind its promoter.

### Bile salts and N-acetyl-D-glucosamine induce native *vieSAB* expression

Because the *vieSAB* operon in the El Tor biotype is poorly expressed during growth in LB, we sought to identify culture conditions or compounds which gave rise to expression of *vieSAB* from its native locus. Identifying such conditions or signals not only allows study of the *vieSAB* operon in its natural context instead of during artificial overexpression, but also provides insight into the environmental conditions in which *V. cholerae* El Tor might benefit from *vieSAB* expression.

We started with a screen for growth conditions or media additives that resulted in expression from the native chromosomal *vieSAB* locus. As a reporter for transcription we used an El Tor *V. cholerae* strain in which promoterless genes for ampicillin resistance (ApR) and kanamycin resistance (KmR), are inserted directly downstream of the *vieSAB* operon such that transcriptional read-through of *vieSAB* would confer resistance to Ap and Km. We screened for inducing conditions that allowed for survival and growth in elevated Ap and Km concentrations as determined by enumeration of CFU after the challenge (Fig 2A-B).

**Fig 2.**
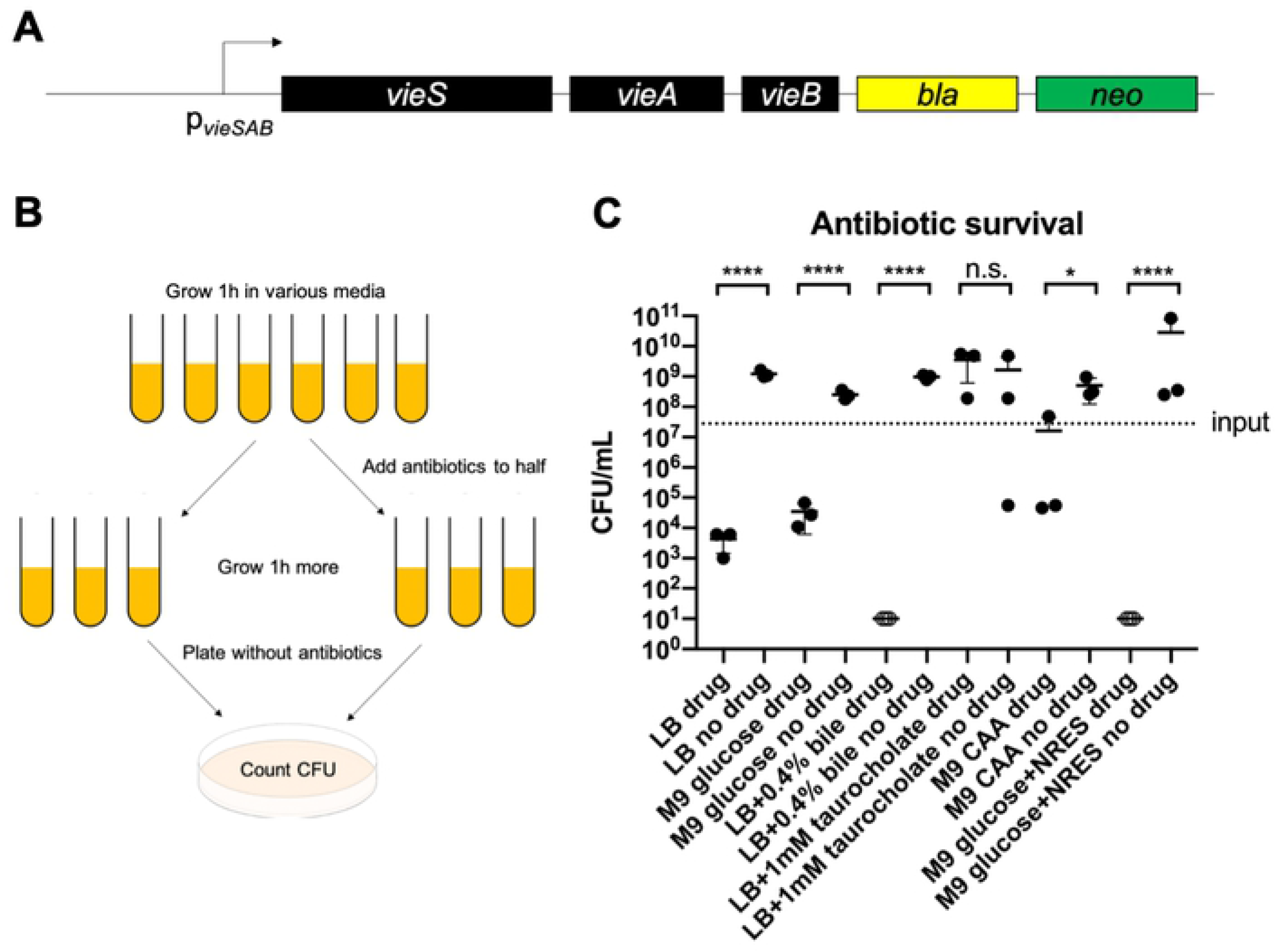
Screen for activators of the *vieSAB* operon. Survival of strains harboring a *vieSAB* transcriptional fusion to dual antibiotic resistance genes at antibiotic concentrations that kill the uninduced fusion strain. A) Genomic context of antibiotic resistance reporters. B) Experimental setup for *vieSAB* inducer screen. C) Survival after antibiotic challenge in the indicated conditions.

Conditions tested included those listed in Table 3, all of which we chose because they are physiologically relevant and could plausibly act as signals of entry into the human host from the aquatic environment. During stationary phase, wild-type *V. cholerae* is much less sensitive to antibiotics than during exponential growth. Because of this phenotype, we were unable to satisfactorily screen the effect of conditions which required growth to stationary phase, including growth in AKI medium and biofilm growth on chitin flakes.

**Table 3:** Conditions tested for *vieSAB* induction.

|  |  |
| --- | --- |
| LB medium | No additives—LB only |
|  | 0.4% porcine bile |
|  | 1 mM cholate (primary bile acid) |
|  | 1 mM taurocholate (secondary bile acid) |
|  | 1 mM deoxycholate (secondary bile acid) |
| | 100 μM ferrous sulfate (iron source) |
|  | 1 mM spermine NONOate (medium half-life nitric oxide donor; nitric oxide is produced by immune cells) |
|  | 0 g/L sodium chloride |
|  | 20 g/L sodium chloride |
|  | 40 g/L sodium chloride |
| M9 medium | Glucose |
|  | Pyruvate |
|  | N-acetyl-D-glucosamine (GlcNAc) |
|  | Casamino acids (CAA) |
|  | Porcine mucin |
|  | Glucose + 12.5 mM/ea L-asparagine, L-arginine, L-glutamic acid, and L-serine |

Initial screening conditions are presented in Fig 2C; remaining conditions were tested in subsequent assays but none conferred increased survival to antibiotics. As expected, growth in the negative control conditions LB and M9 glucose did not confer antibiotic resistance via read-through of the *vieSAB* operon. Whole bile had the opposite effect of sensitizing the cells to antibiotics. However, individual bile acids such as taurocholate, significantly improved survival of the cultures. For alternate carbon sources, casamino acids gave a modest improvement of survival that was not significant upon further study. We also tested growth in M9 glucose supplemented with the four amino acids N, R, E and S, which were previously hypothesized to be a periplasmic signal detected by VieS (Martinez-Wilson et al., 2008). The addition of N, R, E and S, like whole bile, appeared to sensitize the cells to antibiotics; this suggests that these amino acids do not induce expression of the *vieSAB* operon.

To more quantitatively measure *vieSAB* expression in the inducing conditions we identified, we repurposed the *V. cholerae* strain harboring pBBR-lux with luciferase under the control of P_*vieSAB*_. We grew this strain in the presence of taurocholate or its precursor molecule cholate. We also expanded the carbon sources to include pyruvate and N-acetyl-D-glucosamine (GlcNAc) which is abundant in the small intestine. We monitored OD_600_ and luminescence in RLU over several hours and calculated relative light units at mid-exponential phase. Growth in the presence of cholate (CC), taurocholate (TC), and on GlcNAc as a carbon source lead to greater light production than the other growth conditions tested (Fig 3).

**Fig 3.**
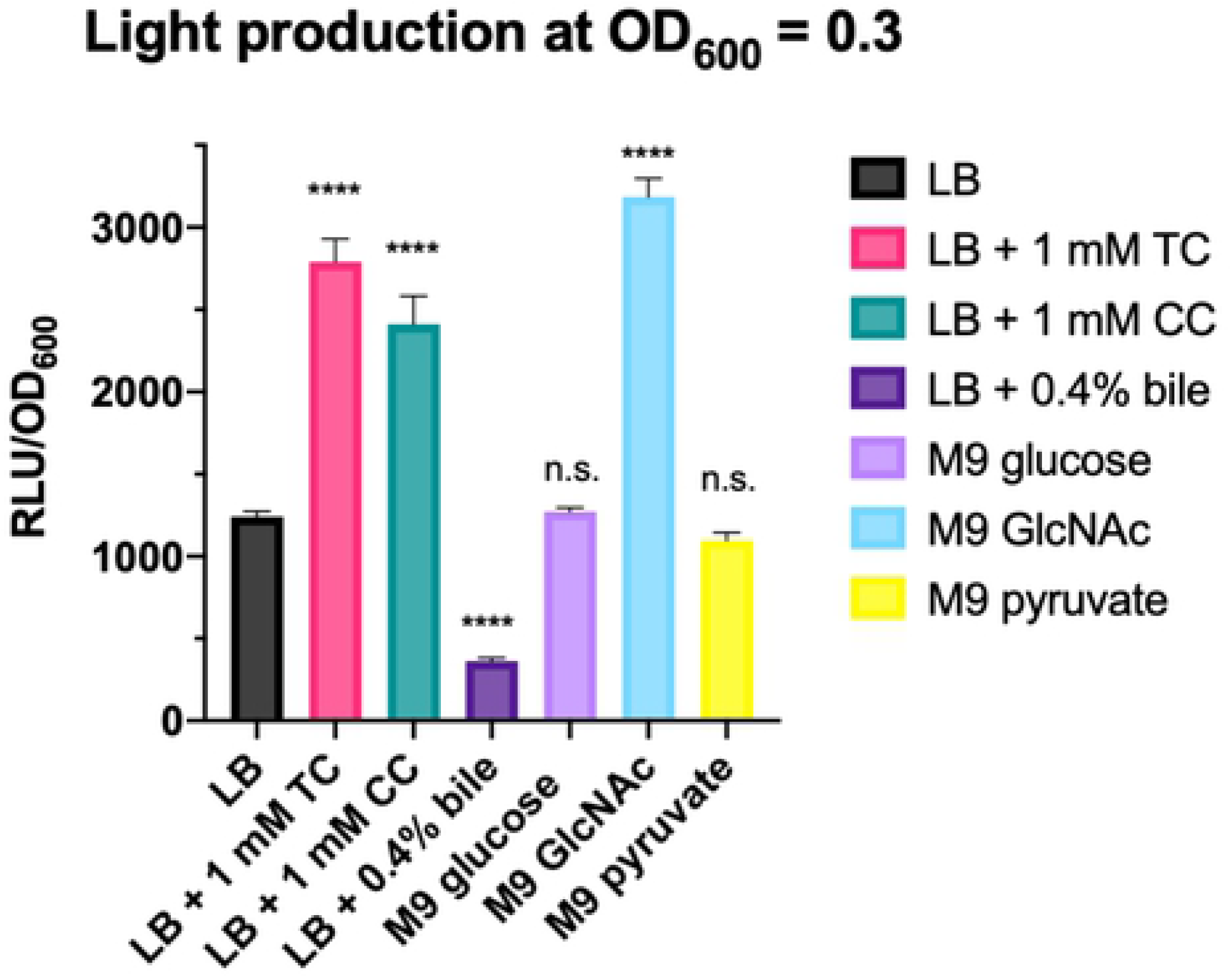
Expression of *vieSAB* in the presence of different activators. Light production driven by the *vieSAB* promoter fused to luciferase was measured after growth to mid-exponential phase (OD_600_ = 0.3) in the listed conditions. (Dunnett multiple comparisons test to LB).

The inducing molecules cholate, taurocholate, and GlcNAc are all present at high concentrations in the duodenum and proximal ileum, the earliest sites of *V. cholerae* colonization after passing through the stomach (Deest et al., 1968; Ghosh et al., 2011). Therefore, these may serve as signals of transition into a human host, triggering expression of the *vieSAB* operon to break down cyclic-di-GMP, in turn leading to virulence-factor expression and motility and repression of biofilm formation. However, GlcNAc is the monomer that makes up chitin, the primary carbon and nitrogen source for *V. cholerae* in the aquatic environment. We wanted to test whether *V. cholerae* can distinguish between monomeric GlcNAc and polymeric chitin; if *V. cholerae* uses GlcNAc as a signal of host colonization it needs to distinguish monomeric GlcNAc from chitin. A difficulty in testing this hypothesis is the fact that chitin is insoluble in water. In order to directly compare growth on GlcNAc to growth on chitin by luciferase assay, we short oligomers of GlcNAc that are slightly soluble in water and have been shown to induce natural competence in *V. cholerae* much like chitin, while monomeric GlcNAc cannot (Meibom et al., 2005). We chose a GlcNAc pentamer, penta-N-acetylchitopentaose ([GlcNAc]_5_) (Seikagaku Biobusiness), as a soluble stand-in for chitin to test expression of *vieSAB*. The solubility of [GlcNAc]_5_ in water is approximately 1 mg/mL or 0.1% w/v, so we modified M9 minimal medium to contain 0.1% w/v of the chosen carbon source instead of the typical 0.4%. This modification allows us to directly compare luciferase expression during growth on glucose, GlcNAc, or [GlcNAc]_5_. We also added 1 mM cholate, 1 mM taurocholate, or 500 μM each cholate and taurocholate to the media to examine the combined effects of bile salts and GlcNAc on *vieSAB* expression. The lower amounts of carbon source used in this experiment resulted in limited growth of *V. cholerae*, and therefore we had to measure light production at a lower culture density than in the previous experiment. While monomeric GlcNAc results in higher *vieSAB* expression than glucose, [GlcNAc]_5_ does not (Fig 4), suggesting that *V. cholerae* can indeed distinguish between monomeric GlcNAc and chitin, and that GlcNAc induces *vieSAB* expression while chitin does not. When [GlcNAc]_5_ is combined with cholate, taurocholate, or both, light production increases though not to the same levels as during growth on GlcNAc.

**Fig 4.**
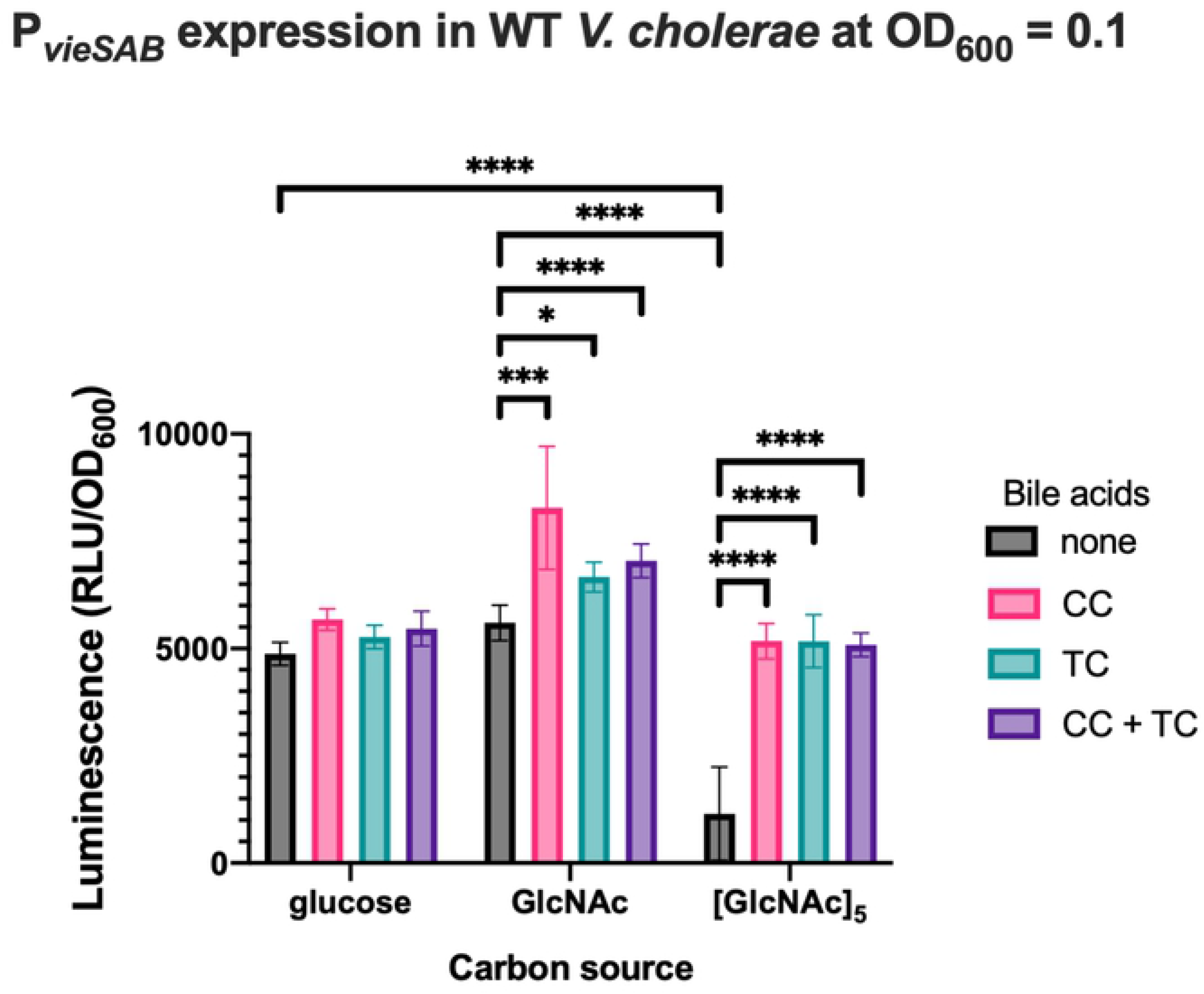
Expression of *vieSAB* responds to N-acetyl-D-glucosamine not soluble chitin. Expression of the *vieSAB* promoter fused to luciferase in modified M9 medium in each indicated condition at mid-exponential growth phase (OD_600_ = 0.1). (Dunnett multiple comparisons test).

## Discussion

The role of cyclic-di-GMP as a signaling molecule in classical biotype *V. cholerae* has been well-studied, but c-di-GMP signaling in El Tor biotype strains is less defined. While we focused exclusively on VieA, it also became clear that the El Tor isolate we studied is less responsive to changes in the overall concentration of c-di-GMP than classical biotype strains; when VieA is deleted or catalytically inactive in classical biotype strains, phenotypes such as reduced motility, increased biofilm formation, and a virulence defect are apparent (Tischler and Camilli, 2005; Martinez-Wilson et al., 2008). However, when the same deletion is made in a wave three El Tor biotype strain, no such phenotypes can be observed (Hammer and Bassler, 2009; Beyhan et al., 2006; Tamayo et al., 2008). Our work on this specific facet of *V. cholerae* biology revealed that phenotypes relating to virulence and colonization are actively evolving as new waves of the pandemic appear.

We examined how expression of the *vieSAB* operon is regulated, revealing that *V. cholerae* activates *vieSAB* transcription upon sensing bile acids and GlcNAc, signals that we believe represent a transition from the aquatic environment to the human host. Bile is present at high concentrations in the portion of the small intestine nearest the stomach, where *V. cholerae* must begin to establish colonization in order to replicate and spread throughout the entire organ. The presence of bile is an indicator of having passed through the harsh acidic environment of the stomach and into the small intestine. The regulatory effects of bile on *V. cholerae* are complex and at times seem contradictory. Although the fatty-acid components of bile inhibit ToxT activity and some bile acids enhance biofilm formation, bile acids also increase cholera toxin expression in a ToxT-independent manner and taurocholate has been shown to activate the ToxR regulon by enhancing TcpP activity (Plecha and Withey, 2015; Hung et al., 2005; Hung and Mekalanos, 2005; Yang et al., 2013). In order to establish an infection *V. cholerae* must express virulence factors, increase its motility, and suppress biofilm formation—all behaviors that are associated with low c-di-GMP concentration. This led to our model that the activators of *vieSAB* expression serve as signals of infection initiation, leading to expression of VieA to break down c-di-GMP and favor virulence-related phenotypes. This model was strengthened by our exploration of carbon source availability as an activator of *vieSAB* expression.

Because chitin is insoluble in water and our screen relied on growth in liquid media, we substituted GlcNAc for chitin in the belief that since *V. cholerae* must break down chitin into GlcNAc prior to metabolism, GlcNAc would trigger the same behavioral changes in *V. cholerae* as chitin does. With this context in mind, we were surprised to find that growth in minimal media with GlcNAc as the sole carbon source led to increased *vieSAB* expression. Monomeric GlcNAc is very abundant in the small intestine (Deest et al., 1968; Ghosh et al., 2011), and therefore if *V. cholerae* can distinguish between monomeric GlcNAc and chitin, then an abundance of the monomer and an absence of chitin could serve as a signal of environmental transition. This is not unprecedented. There is evidence that *V. cholerae* distinguishes between monomeric and oligo- or polymeric GlcNAc or chitin; for example, GlcNAc does not induce natural competence in the El Tor biotype while chitin does. We found a soluble compound consisting of a GlcNAc pentamer, penta-N-acetylchitopentaose, which does induce natural competence and thus which we believe is a suitable soluble stand-in for chitin in contrast to monomeric GlcNAc (Meibom et al., 2005). We then found that growth in minimal media with GlcNAc as a carbon source induced *vieSAB* expression, while growth in the same minimal media with penta-N- acetylchitopentaose does not. We do not yet know the mechanism by which *V. cholerae* distinguishes between GlcNAc and chitin, but our finding that only GlcNAc activates *vieSAB* transcription combined with the knowledge that only chitin activates natural competence shows that the distinction is indeed made. The presence of bile salts and GlcNAc at the same time is a near-unmistakable signal; this combination is only present when *V. cholerae* has reached the small intestine of the host.

Our overall model is that *V. cholerae* senses GlcNAc, cholate, and taurocholate as signals of entry into the small intestine, expression of *vieSAB* increases because VieA is beneficial though nonessential during early stages of infection to establish initial colonization. This aspect of our model is also consistent with the suggestion that cyclic-di-GMP must be depleted early during infection to enhance colonization of the El Tor biotype (Tamayo et al., 2008) Later in the infection, *V. cholerae* must prepare for reentry into the aquatic environment. As cell density increases during late stages of infection, quorum sensing comes into play and HapR expression increases. The *vieSAB* operon is silenced by HapR, which we propose is a part of the preparation for aquatic survival; the cells no longer need to express virulence factors or retain high motility, so VieA expression is halted which will then favor an increase in intracellular c-di-GMP and a reversion to behaviors such as biofilm formation.

## Acknowledgments

We thank members of the Camilli lab for helpful comments on this work, especially David Lazinski, Neil Greene, and Jacob Bourgeois. We thank Rebecca Zhang for carrying out preliminary assays for the screen for activators of *vieSAB*.

